# Macrophages Enforce the Blood Nerve Barrier

**DOI:** 10.1101/493494

**Authors:** Liza Malong, Ilaria Napoli, Ian J White, Salome Stierli, Alessandro Bossio, Alison C. Lloyd

**Affiliations:** MRC Laboratory for Molecular Cell Biology, University College London, Gower Street, London, WC1E 6BT, UK

**Author notes:** These authors contributed equally to this manuscript (Liza Malong, Ilaria Napoli).

## Abstract

The specialised blood barriers of the nervous system are important for protecting the neural environment but can hinder therapeutic accessibility^1,2^. Studies in the central nervous system (CNS) have shown the importance of the cellular components of the neuro-vascular unit for blood-brain barrier (BBB) function. Whilst the endothelial cells (ECs) confer barrier function with specialised tight junctions (TJs) and low levels of transcytosis, pericytes and astrocytes provide complete coverage of the ECs and both deliver essential signals for the development and maintenance of the BBB^3–9^. In contrast, the blood-nerve barrier (BNB) of the peripheral nervous system (PNS) remains poorly defined^10^. Here, we show that the vascular unit in the PNS has a distinct cellular composition with only partial coverage of the BNB-forming ECs. Using a mouse model, in which barrier function can be controlled^11^, we show the BNB, while less tight than the BBB, is maintained by low levels of transcytosis and the TJs of the ECs, with opening of the barrier associated with increased transcytosis. Importantly, we find that while ECs of the PNS have higher transcytosis rates than those of the CNS, the barrier is reinforced by resident macrophages that specifically engulf leaked material. This identifies a distinct role for macrophages as an important component of the BNB acting to protect the PNS environment with implications for improving therapeutic delivery to this tissue.

---

To investigate the cellular structures comprising the BNB in the PNS, we took complementary approaches. To determine the total cellular coverage of the ECs in peripheral nerve in a non-biased fashion, we performed 3D EM of endoneurial blood vessels that can detect all cellular contacts (Fig. 1a, Extended Data Fig. 1a, Movie1). In contrast to the CNS^12^, we found cellular coverage of PNS endoneurial blood vessels was incomplete, a finding confirmed by higher-resolution 2D EM images (Fig1b and Extended Data Fig. 1b). Pericytes could be identified by their tight association with the ECs within the basal lamina, however, other cell types were observed, which made looser contacts with the ECs. In a separate study, we have used immunostaining to identify all cell-types within the endoneurium of peripheral nerve^13^ and we used these markers and others, to determine the cell-types that interact with ECs in the PNS and verified these findings using CLEM. We found 3 different populations of cells that directly interact with endoneurial blood vessels. First, classical pericytes, expressing the markers NG2, PDGFRβ and α-SMA provided partial coverage of the ECs and were embedded in their basement membrane (Fig. 1c, Extended Data Fig. 2). The percentage of pericyte coverage varied greatly between vessels (10-100%, with a mean of 54%), despite all vessels having barrier function (Figure 1d, Extended Data Fig. 2). Second, a population of fibroblasts/pericytes-like cells that express NG2, PDGFRβ, p75 and CD34 were found more loosely associated with the ECs, outside of the basement membrane. These cells displayed a characteristic, elongated morphology and made multiple contacts with the basement membrane of the vessels (Fig.1c,e, Extended Data Fig. 3a-d). Finally, we found that a population of macrophages that express Iba1 and F4/80 was intimately associated along the vessels (Fig.1c,e, Extended Data Fig. 3c-e)^13^. These findings demonstrate that the vascular unit that comprises the BNB is distinct from the neurovascular unit of the BBB, consisting of ECs partially covered by pericytes with more loosely associated pericyte-like cells and macrophages (Extended Data Fig.4a).

**Figure 1.**
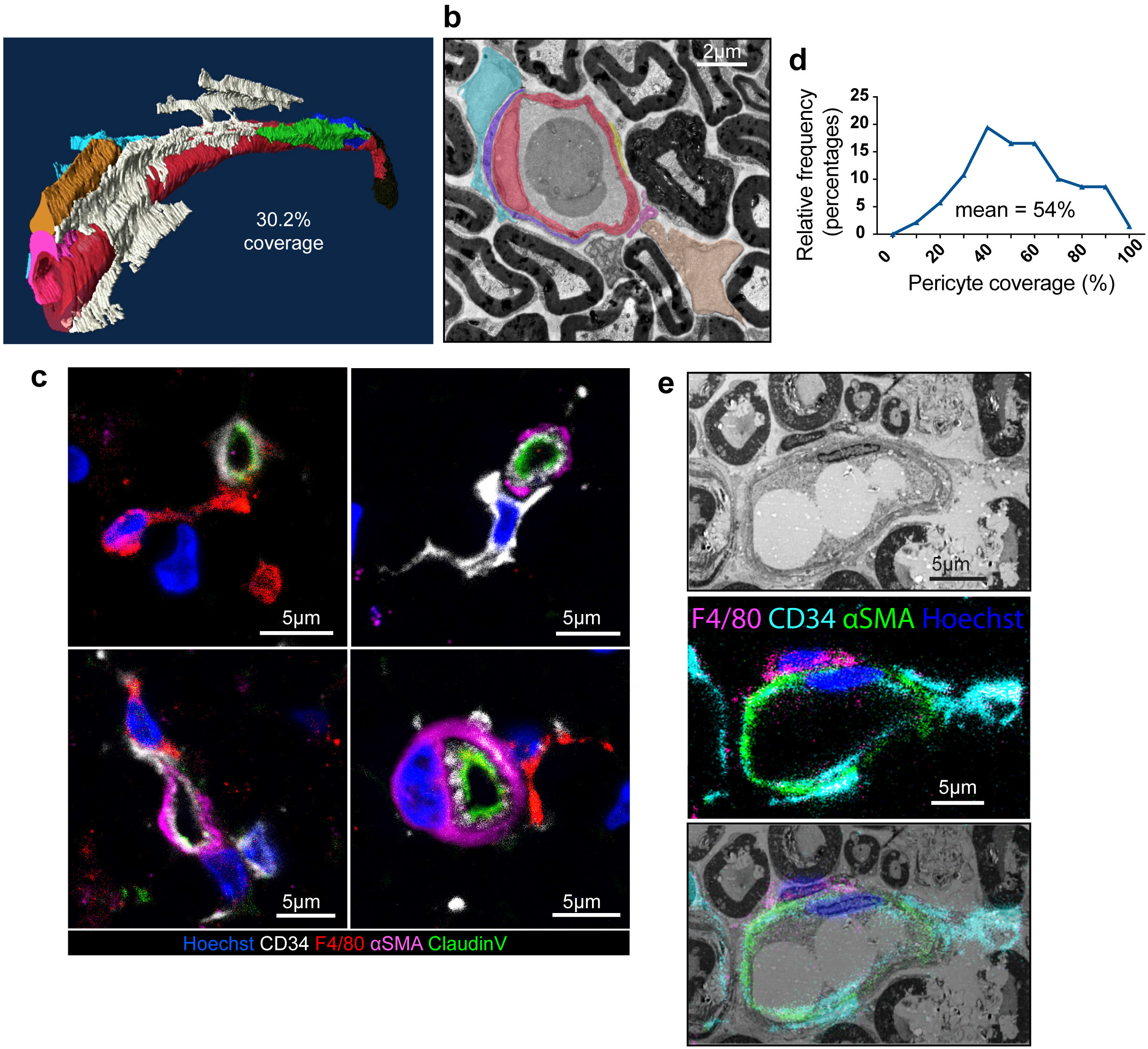
Structure of the BNB. **a**, 3D EM reconstruction of an endoneurial blood vessel and contacting cells. **b**, Representative transverse EM image of the vascular unit. ECs are coloured red, whilst other cell types associated with the blood vessel are individually coloured. **c**, Representative fluorescent images of the vascular unit in the endoneurium. ECs (CD34, white and claudin V, green), pericytes (αSMA-magenta), pericyte-like cells (CD34, white) and macrophages (F4/80, red), nuclei (Hoechst, blue). Images show the variable coverage of the vessels with pericytes and the close interactions of pericyte-like cells and macrophages with the vessels. **d**, Quantification of pericyte coverage visualised by EM. Graphs shows the frequency distribution. (n= 6 animals, 139 blood vessels). **e**, Representative CLEM images of the BNB vascular unit. ECs and pericyte-like cells (CD34, cyan), macrophages (F4/80, magenta), pericytes (αSMA, green), nuclei (Hoechst, blue).

In the CNS, endothelial barrier function is associated with specialised TJs and with extremely low levels of transcytosis^5,6,14–17^. To determine the nature of the barrier of the PNS, we injected Horseradish Peroxidase (HRP) into the tail veins of mice and used EM to determine the permeability of the TJs and the levels of transcytosis. Importantly, we compared blood vessels within the endoneurium, which have barrier function, to those between the nerve fascicles, which do not (Extended Data Fig. 4b-c)^18^. Comparison of the ECs of the epineurial and endoneurial blood vessels showed they had differences both in the structures of their tight junctions and their rates of transcytosis. The tight junctions of the epineurial ECs (EpiECs) were penetrated to varying degrees by the HRP, with fewer than 20% showing zero penetrance. In contrast, the majority of endoneurial ECs (EndoECs) were not penetrated with HRP with a “kissing point” very close to the inner surface of the ECs (Fig. 2a-b).

**Figure 2.**
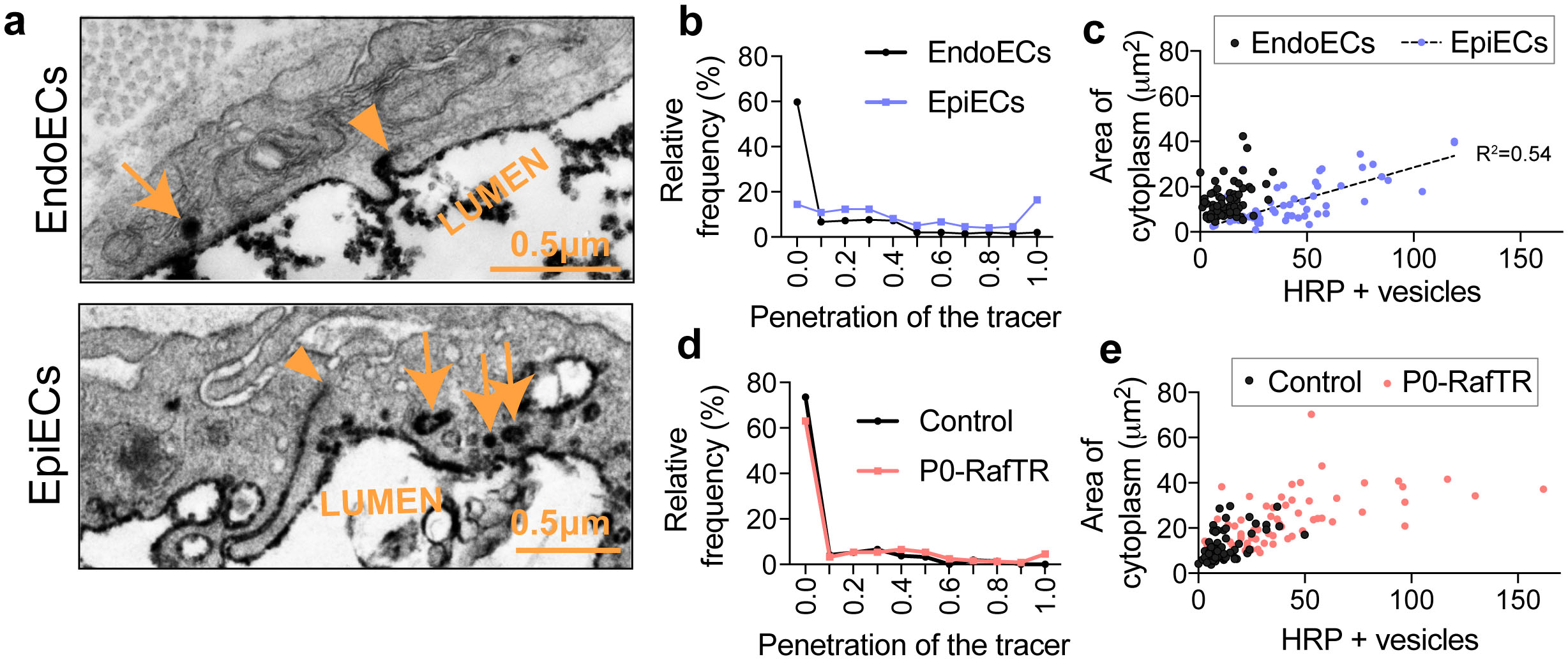
Regulation of the BNB. **a**, Representative EM images of endoneurial and epineurial blood vessels from mice injected with HRP. HRP+ vesicles are marked by arrows and penetration of HRP into the junctions is marked by arrowheads. **b**, Graph shows the relative frequency of the penetration of HRP into the junctions between endoneurial (black) and epineurial (blue) ECs following IV injection of HRP (n= 3 animals, 55/52 blood vessels and 194/194 junctions). **c**, Graph shows the area of cytoplasm of ECs against the number of HRP+ vesicles. Each dot represents an individual blood vessel. **d**, Relative frequency of the penetration of the HRP into the junctions between endoneurial endothelial cells from Control (black) and P0-RafTR (red) animals, 10 days after the first tamoxifen injection. (n= 3/4 animals, 79/45 blood vessels and 212/240 junctions). **e**, Graph shows the area of cytoplasm of ECs against the number of HRP+ vesicles in tamoxifen treated control and P0-RafTR animals at Day 10. Each dot represents an individual blood vessel.

Likewise, the levels of transcytosis were different between the barrier and non-barrier blood vessels within the nerve with the EpiECs containing a considerably higher numbers of vesicles and a higher vesicular density compared to the EndoECs (Fig 2a, c and Extended Data Fig. 4d-e). Interestingly, the differences in vesicular density mirror those found between immature retinal blood vessels (no barrier function) and the mature blood vessels that constitute the blood retinal barrier^14^. Notably, the vesicular density was slightly higher in the EndoECs compared to those in the mature vessels of the retina, which is consistent with the more permeable nature of the BNB compared to those of the CNS^14^.

The BNB is regulatable in that following an injury to the nerve, the barrier opens, coincident with the inflammatory response that is important for the regeneration of peripheral nerve^11,19^. Using a transgenic mouse model in which the ERK-signalling pathway can be reversibly induced specifically in myelinating Schwann cells (mSCs) (P0-RafTR mice), we have shown previously that the barrier can be opened by signals downstream of ERK signalling in SCs (Extended Data Fig. 4f)^11^. This control of the barrier is part of the process by which dedifferentiated adult Schwann cells orchestrate the regenerative response but importantly occurs in the absence of trauma or axonal damage, providing a powerful mouse model for studying the regulation of the BNB.

To further explore the nature of the BNB, we determined how it can be “opened” in the P0-RafTR mouse model. We first analysed the perineurium, which provides an external barrier to the nerve fascicles. EM analysis of nerves at Day 10 following the first tamoxifen injection, when the BNB is fully opened, showed minimal structural changes. Moreover, immunostaining with antibodies to the junctional proteins VE-Cadherin and Z01 showed the junctions between the perineurial cells appeared unchanged and the integrity of the perineurium remained intact, as determined by the failure of 40kD dextran to permeate the perineurium either *in situ* or *ex-vivo* (Extended Data Fig. 5–6).

We next analysed the blood vessels of the endoneurium. EM analysis showed that the blood vessels looked different in P0-RafTR compared to control Tmx-treated animals in that they appeared less regular and compact in shape, with a larger cytoplasmic area and significantly more protrusions into the lumen (Extended Data Fig. 7a-c). To analyse the functionality of the TJs, we injected the mice with HRP, but did not find any significant differences in the tightness of the junctions between endothelial cells (Figs 2d and Extended Data Fig.7d). Consistent with this, analysis of VE cadherin and ZO1 did not reveal any structural differences in the EC junctions implying that changes in the TJs are not responsible for the opening of the BNB (Extended Data Fig.7e).

Importantly however, we found a dramatic increase in the transcytosis rates in the P0-RafTR mice, with an increase in both the number of HRP filled vesicles per vessel and an increase in vesicular density to levels seen in the non-barrier vessels of the epineurium (Fig. 2e, Extended Data Fig. 8a-c). Furthermore, this finding was confirmed with the use of a second tracer, BSA-FITC (Extended Data Fig 8d-e). These results show that the opening of the BNB is independent of changes in perineurial or TJ permeability but are instead the result of increased transcytosis levels in EndoECs.

Analysis of the vascular unit in the P0-RafTR mice showed that the level of pericyte coverage remained unchanged when the barrier was open, despite the increase in transcytosis rates (Extended Data Fig. 9a-b). Moreover, the pericyte-like cells remained associated with the blood vessels (Extended Data Fig 9c-d). However, we did detect a dramatic change in the perivascular localisation of the macrophages in that while there was a large increase in the number of macrophages within the nerve, coincident with the opening of the BNB, there were far fewer associated with the blood vessels (Extended Data Fig. 9e-f, Movie 2).

The perivascular localisation of macrophages indicated a potential role in barrier function. The BNB is more permeable than the BBB and we observed in animals injected with HRP that the low levels of transcytosed HRP rapidly (<1 minute) accumulated in endoneurial macrophages. The HRP accumulated in a time-dependent fashion and by 6 hours, the macrophages were filled with HRP (Fig 3a). Strikingly, the two other cell populations that interact with the ECs, pericytes and the pericyte-like population failed to accumulate any detectable HRP even at the 6 hour time-point (Fig 3b-c). This remarkably efficient “vacuuming” of material that enters the endoneurium suggested that while the higher levels of transcytosis of the EndoECs make the BNB more permeable than the BBB, this may be may be compensated for by macrophages.

**Figure 3.**
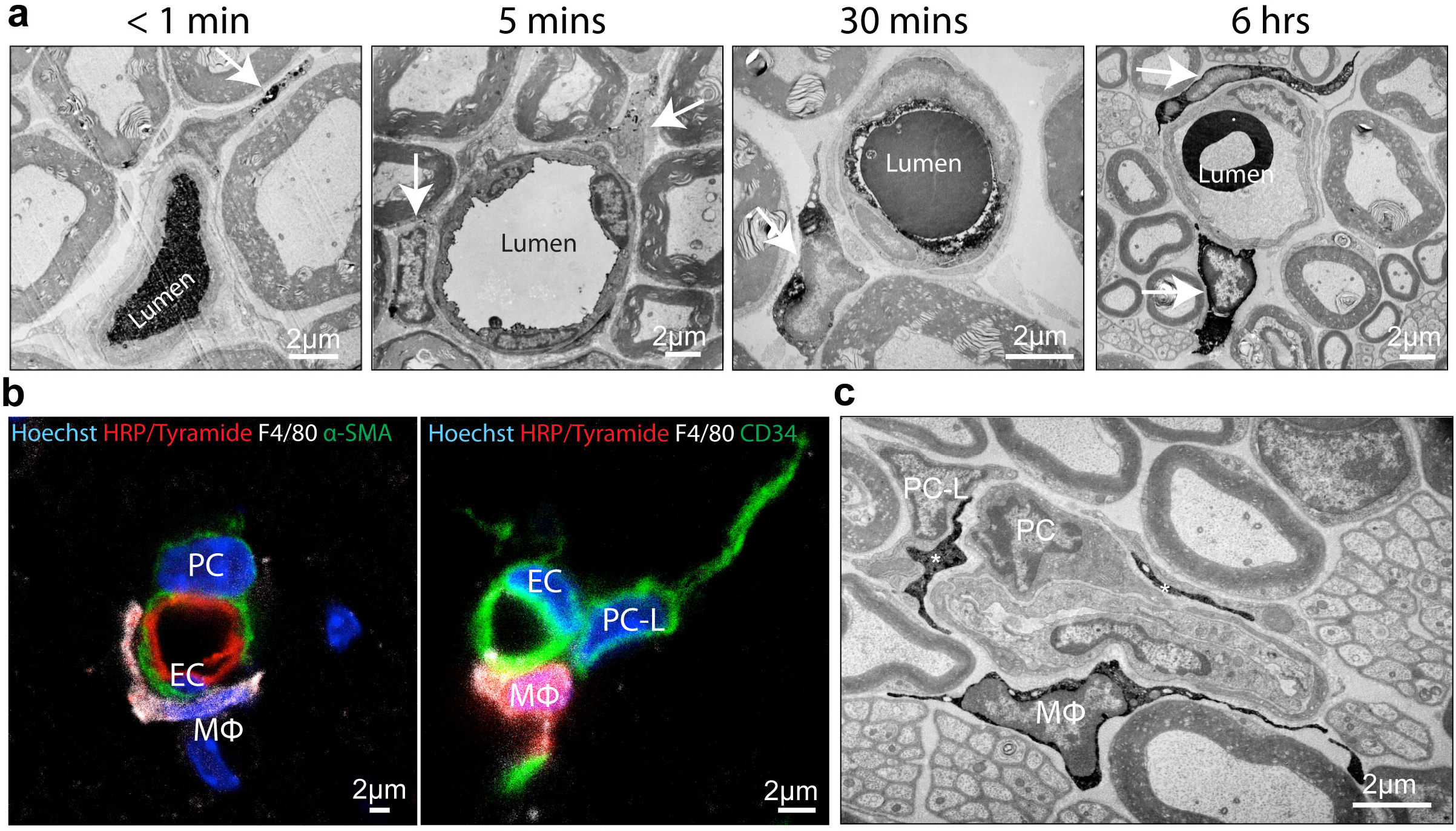
Macrophages specifically and rapidly engulf material that leaks through the BNB. **a**, Representative EM images showing that macrophages accumulate HRP. Animals were injected IV with HRP and harvested immediately (<1min), 5 mins, 30 mins and 6 hrs later. Arrows point to perivascular macrophages containing HRP (n=3 animals at each time point). **b**, Fluorescent confocal images showing that whereas macrophages (MΦ, white) take up HRP (red), pericytes ((PC) left, green) and pericytes-like cells ((PC-L) right, green) do not. Nuclei are stained with Hoechst (blue). **c**, Representative EM image showing that pericytes and pericytes-like cells do not accumulate HRP, as opposed to macrophages marked by Mφ and asterisks. Animals were harvested 6 hrs after HRP IV injection for **(b)** and **(c)**.

To test this hypothesis, we used the drug PLX5622 to remove macrophages from peripheral nerves. This small compound drug targets CSF1-R, a tyrosine kinase receptor required for macrophage survival and proliferation and has been shown to efficiently deplete macrophages and microglia *in vivo* in the nervous system^20^’^21^. We found that administration of PLX5622 for 11-12 days led to an almost complete depletion of macrophages in peripheral nerves, with no detectable effects on the number or structure of blood vessels or the other interacting cell types (Figs 4a, Extended Data Fig. 10a-c). Importantly, the permeability of the TJs and the transcytosis rates remained unchanged showing that macrophages were not controlling the tightness of the EC barrier (Figs. 4b-d, Extended Data Fig. 10d-e). However, following HRP injection into the circulation, a dramatic accumulation of HRP could be observed throughout the endoneurium. Remarkably, this leakage could be observed at the gross level of the nerve following reaction with DAB (Fig.4e). Fluorescence microscopy showed that in contrast to control nerves, where HRP was detectable only within the blood vessels and macrophages, nerves lacking macrophages showed high levels of HRP throughout the endoneurium (Fig. 4f-g). Quantification showed the area of HRP staining was 23.82±8.05% of the endoneurium (Fig. 4g), corresponding to the known 20-25% intrafascicular volume^18^ consistent with the spread of leaked material throughout the endoneurium of the nerve. EM analysis further confirmed this finding with HRP detected in the spaces between mSCS in regions obscuring the collagen bundles seen in control nerves (Fig. 4h). These findings demonstrate that macrophages provide a secondary barrier to enforce the protection provided by the ECs and this mechanism is essential to maintain the nerve environment free from blood-borne molecules.

**Figure 4.**
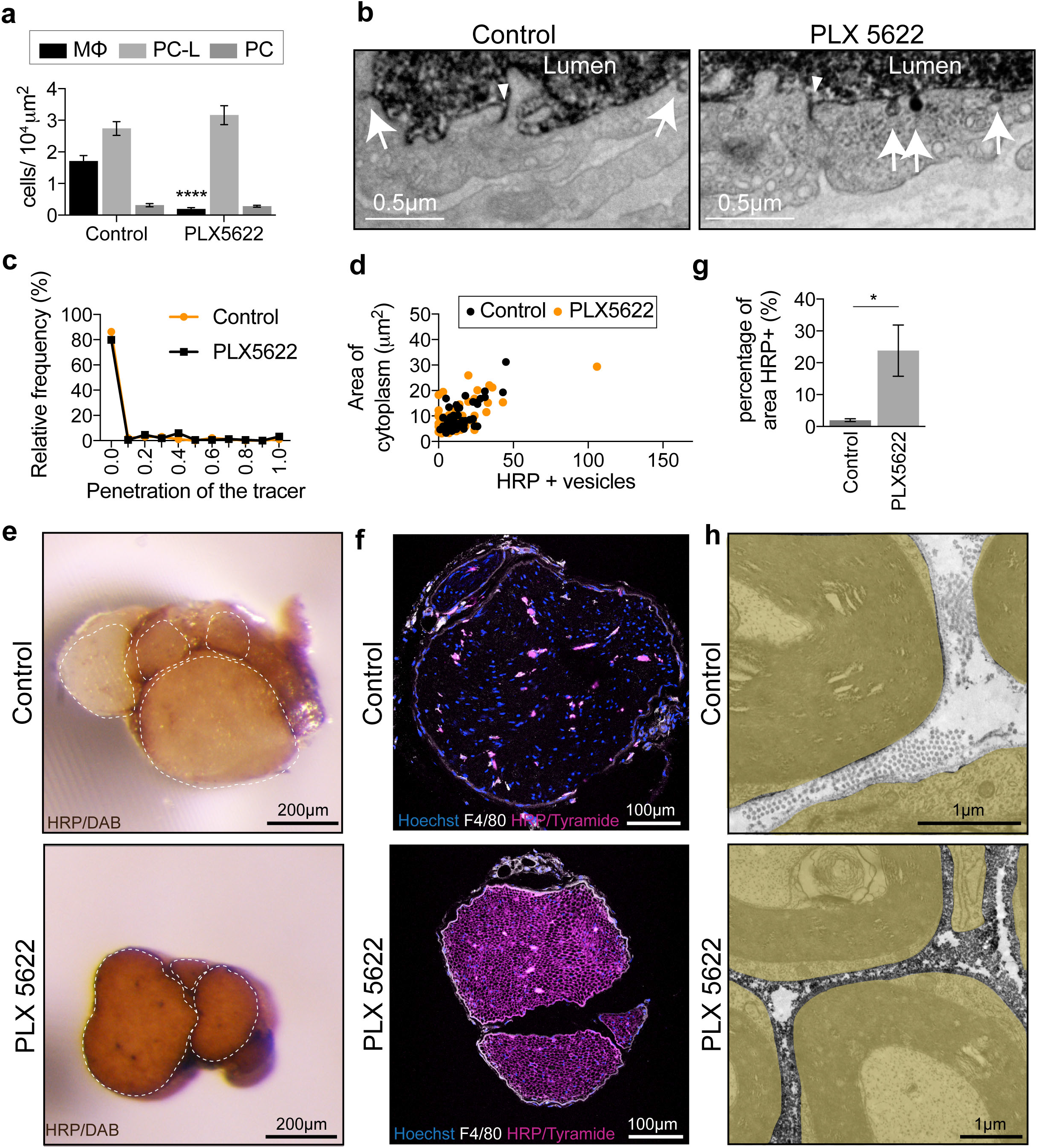
Macrophages enforce the BNB. **a**, Quantification of the number of macrophages (F4/80+/Iba1+), pericyte-like cells (CD34+) and pericytes (αSMA+) in the endoneurium of animals treated with Control or PLX5622 containing chow (mean±SEM, n=14/16 animals for macrophages and 7/9 animals for pericytes and pericytes-like cells respectively). **b**, Nerves from Control (orange) or PLX5622 (black) treated animals were harvested 5 minutes after IV injection of HRP. Representative EM images of the EndoECs with HRP+ vesicles indicated by an arrow and the penetration of the junctions marked by an arrowhead. Quantification of **(b)** showing **c**, Relative frequency of the penetration of HRP into the junctions between EndoECs, **d**, Area of cytoplasm of ECs against the number of HRP+ vesicles. Each dot represents an individual blood vessel. **e**, Representative images of 150μm thick nerve sections stained with DAB. **f**, Representative fluorescent images of tyramide reactivity (magenta) in the endoneurium of control and PLX5622 treated animals. **g**, Quantification of **(f)** showing the percentage area of HRP reactivity in the endoneurium of Control and PLX5622 treated animals (n= 5/6 animals). **h**, Representative EM images highlighting the accumulation of HRP in the extracellular space of the endoneurium in PLX5622 treated animals. Cells are coloured brown.

Here, we have defined the vascular unit that confers the barrier function of the PNS and show that it is distinct from the BBB both in the degree of coverage and the cell-types that comprise the unit. Furthermore, we show that the low transcytosis rates that confer barrier function allows “leakage” into the endoneurium and identify a critical role for macrophages as a second line of defence to enforce the BNB. These findings have important implications for our understanding of this important vascular defence system that guards the PNS and suggests new approaches for both increasing the protection and the delivery of therapeutics to this tissue.

## Methods

### Mouse strains

The P0-RafTR mouse strain was produced by us and is available at The Jackson Laboratory (B6;CBA-Tg(Mpz-RAF1/ESR1)A668Ayld/J)^11^. P0-RafTR animals express an inducible Raf kinase/estrogen receptor fusion protein under the control of the P0 promoter. During adulthood, expression is restricted to myelinating Schwann cells in the PNS. The NG2dsRED mouse strain (NG2-dsRedBAC, IMSR_JAX:008241)^22^ expresses the DsRed.T1 protein under the control of the NG2 promoter.

### Administration of substances

Tamoxifen (Sigma, T5648) was dissolved in EtOH at a concentration of 200mg/ml and further diluted in sunflower oil (Sigma, S5007) to reach a final concentration of 20mg/ml. The solution was vortexed for >3 hours to ensure total dissolution. Adult P0-RafTR and littermate controls were injected with a daily dose of tamoxifen at 150-200μg/g of body weight intraperitoneally for 5 days and harvested at Day 10.

PLX5622 was formulated in AIN-76A chow at 1200ppm and kindly provided by Plexxikon Inc. Animals were provided free access to either PLX5622 containing chow or control chow for 11-12 days before harvesting.

Intravenous tracers were injected in the tail vein of adult animals. Evans blue solution was freshly prepared by dissolving Evans Blue (Sigma, E2129) and BSA (Sigma, A3294) in PBS to reach a concentration of 1% and 5%, respectively. Each animal was injected with 10μl/g of body weight and harvested 30 minutes later. HRP solution was prepared by dissolving 125mg of HRP type II (Sigma, P8250) in 2.5ml PBS. Each animal was injected a single dose of 0.5mg/g of body weight and were harvested at various time points. Lysine fixable BSA-FITC (Invitrogen, A23015) was prepared at 1.5 mg/ml in PBS. Animals were injected at 10μl/g of body weight and were harvested 30 minutes later.

### Immunostaining

Immunostaining was performed as described in Napoli et al 2012^11^. Briefly, nerves were fixed in 4% paraformaldehyde (PFA) for 4-6 hours at RT, cryopreserved in 30% sucrose O/N at 4°C, and 15% sucrose/50 % OCT for 2 hours at RT, prior to being embedded in OCT and snap frozen. 10μm sections were obtained with a cryostat, permeabilised in 0.3% Triton/PBS for 30 mins at RT, and blocked in 10% goat serum/PBS for 1 hour at RT, prior to the primary antibody incubation. When Claudin V antibody was used, the sections were blocked in 2% Fab fragment (Jackson Immunoresearch 715-007-003) /10% Donkey Serum/PBS O/N at 4°C instead, and donkey serum and secondary antibodies raised in donkey were used in the following steps. Antibodies used were: F4/80 (BioRad MCA497G 1/400), Iba1 (Wako 019-19741, 1/500), CD31 (BD pharmingen 550274, 1/100), CD34 (abcam ab81289, 1/400), αSMA (Sigma F3777 1/1000), Claudin V (Invitrogen 352500, 1/500). Primary antibodies were diluted in blocking solution and incubated O/N at 4°C. The next day, sections were washed in PBS and incubated with the corresponding secondary antibody, raised in goat, conjugated to Alexa Fluor 488/594/647, for 1 hour at RT. This solution also contains Hoechst (1/1000). To reveal the presence of HRP in HRP-injected samples by light microscopy, an additional step was performed after the secondary antibody incubation. Samples were washed in PBS and incubated in an amplification buffer containing 0.1% tyramide reagent and 0.0015% H_2_O_2_ for 5 minutes in the dark at RT (Molecular Probes T-20935). Finally, sections were washed in PBS, rinsed in water and mounted under a glass coverslip with Fluoromount.

Confocal images were acquired with a SPE3 confocal microscope, (Leica TCS SPE, lasers: 405nm, 488nm, 561nm and 635nm), a SPE8 confocal microscope (Leica TCS SPE8 STED 3x, UV and white light laser 470-670nm) or a multiphoton microscope (LSM880, laser lines 405,488,561 and 633, lambda acquisition).

### 3D immunostaining

Junction protein immunostaining of the perineurium and blood vessels used the PACT protocol^23^. Briefly, nerves were fixed in 4% PFA O/N at 4°C. Nerves were then incubated in 4% acrylamide/0.25% VA-044/PBS O/N at 4°C. Nerves were exposed to nitrogen for 5 minutes and left at RT for 3 hours, prior to being washed in PBS for one day at 37°C. All subsequent incubations took place at 37°C in a rotating device. Nerves were then incubated in 8% SDS for 4 days, with a fresh solution of SDS used on the third day. Nerves were washed in 0.01% Sodium Azide /PBS O/N, prior to the primary antibody incubation. Antibodies were diluted in 2% donkey serum/ 0.1% Triton/ 0.01% sodium azide/PBS and incubated for 3 days, with fresh antibody added every day. The antibodies used were: VE-Cadherin (R&D systems AF1002, 1/10) and ZO1 (Invitrogen 61-7300, 1/10). Nerves were washed O/N in PBS and further incubated in the relevant secondary antibodies coupled to donkey Alexa Fluor fluorophores (1/100). Antibodies were diluted in 2% donkey serum/0.1% Triton/0.01% sodium azide/PBS and incubated for 4 days, with fresh antibody added daily for the first 3 days. Nerves were then washed in PBS O/N and incubated in Histodenz (Sigma D2158, 1.3 g/ml), prior to being mounted in Histodenz using a mould (Grace bio-labs 664113) and dental glue (picodent twinsil).

Blood vessel, pericyte and macrophage immunostaining were performed using the iDisco protocol ^24^ Briefly, nerves were harvested and fixed in 4% PFA O/N at 4°C. The next day, nerves were left in PFA for 1h at RT, then washed in PBS and 0.2% Triton/PBS at RT for 1 hour each. Nerves were then incubated in 0.2%Triton /20%DMSO/PBS O/N at 37°C. The next day, the nerves were incubated in 0.1% Triton/0.1% Tween /0.1%(w/v) deoxycholate /0.1% NP40 /PBS O/N at 37°C. The samples were then washed in 0.2% Triton/PBS twice for 1 hour at RT, then incubated in 0.2% Triton /20% DMSO /0.3% glycine (w/v)/PBS O/N at 37°C. The samples were further washed in 0.2%Tween / 0.001% (w/v) heparin/PBS twice for 1 hour at RT, then blocked in 0.2% Triton /10%/o DMSO /6%/o donkey serum/PBS for 8 hours at 4°C. The nerves were then incubated in a blocking solution containing the primary antibodies (to Iba 1, VE cadherin, αSMA and CD31, all 1:50) for 3 days at 37°C. Fresh antibody was added each day during this incubation. After this, nerves were washed in 0.2%Tween /0.001%heparin (w/v)/PBS >8 hours and further incubated with a blocking solution containing the relevant secondary antibodies (1:100) for 3 days. Fresh antibody was added each day during this incubation. After this, nerves were washed in 0.2%Tween /0.001% (w/v) heparin/PBS for 24 hours. Nerves were then cleared as follows: nerves were first dehydrated by a series of incubations in tetrahydrofuran (THF Sigma 186562) starting with 50% THF/50% water O/N, followed by 1 hour in 80% THF and twice for 1 hour in 100% THF. Nerves were then incubated in dichloromethane (DCM, Sigma 270997) until the sample sank. Finally, nerves were incubated in DiBenzyl Ether (DBE, Sigma 108014), and mounted in DBE using a mould and dental glue.

### Transmission Electron Microscopy (TEM)

TEM was performed as described in Cattin et al 2015^25^. Briefly, nerves were fixed in a phosphate buffer solution (0.2M Na_2_HPO_4_: 0.2M NaH_2_PO_4_ in a 4:1 ratio) containing 2% gluteraldehyde and 2% PFA, for 1 to 7 days at 4°C. After 3 washes in phosphate buffer, the nerves were osmicated in 2% Osmium Tetroxide for 2 hours at 4°C. The solution was rinsed off by 3 washes in water. The samples were then negatively stained in 2% Uranyl acetate for 45 mins at 4°C and washed in water. Nerves were dehydrated by a series of incubations in increasing concentrations of ethanol: 5 mins in 25%, 5 mins in 50%, 5 mins in 70%, 10 mins in 90% and 4x10 mins in 100% ethanol. The dehydrated nerves were then put in contact with a propylene oxide solution for 3x10 minutes, ensuring complete removal of residual ethanol in the tissue. Resin was prepared by mixing TAAB 812 (47%), DDSA (18.5%), MNA (32.5%) and DMP30 (2%). Nerves were then incubated for 1 hour in 50% resin-50% propylene oxide and a 100% resin O/N. The resin was refreshed in the morning. Nerves were embedded in resin in the evening and incubated at 60°C O/N to allow the resin to set.

When the animals were injected with HRP, the nerves were embedded in 2.8% low melting point agarose after the fixation, and sections of 150 microns were prepared with a vibratome. Sections were washed in 0.05M Tris/HCl pH7.6 before undergoing DAB (3,3’-Diaminobezidine tetra-HCl) reaction as follows: the DAB reaction solution was obtained by adding 250μl of a 3% DAB solution (TAAB, D008) to 10ml of Tris/HCl buffer and 7μl of hydrogen peroxidase. Sections were incubated in this solution in the dark for 15 minutes and further washed in the Tris buffer. After this, the sections were osmicated, stained, dehydrated and embedded as described above.

Images were acquired with a FEI Tecnai Spirit Bio-twin electron microscope and a Morada G2 camera (Olympus Soft Imaging Solutions).

### Correlative light electron microscopy (CLEM)

CLEM was performed as described in Cattin et al 2015^25^. Briefly, nerves were fixed in 4% PFA at 4°C. The next day, they were washed in ice-cold PBS for 30 min, and embedded in a 2.6% low melting point agarose gel. The nerves were then sectioned with a vibratome, obtaining 100μm cross sections. The sections were blocked in a solution of 10%GS/PBS for 30 minutes, prior to being incubated in a solution of GS/PBS containing the primary antibodies against CD34 and F4/80 mentioned above (1:400) for 1 hour. The sections were washed in PBS and further incubated for 1 hour in a solution of 10% GS/PBS containing secondary antibodies (goat anti rabbit AF647 and goat anti rat AF594, both 1:400), Hoechst (1:1000), and primary antibody against αSMA conjugated to FITC (1:500). All incubations were performed at 4°C. After this incubation, the sections were placed in a glass-bottom petri dish and fixed with 2.6% low melting point agarose. Fluorescence images were acquired with a SPE3 confocal microscope. Following acquisition, the sections were detached from the petri-dish and fixed for a second time in a 0.2M phosphate buffer containing 2.5% PFA and 1.5% gluteraldehyde for 30 minutes at 4°C. The fixative was washed O/N in phosphate buffer at 4°C. The next day, the sections were subjected to the traditional EM protocol as described above. Ultra-thin sections were prepared with a microtome as described above, with particular care of collecting the first, most superficial, sections. EM images were acquired and processed as described above. The correlation between the fluorescent images and the electron microscopy images were performed by hand, using FIJI, Adobe Photoshop and Adobe Illustrator.

### 3D TEM

The 3D TEM was performed as described in Straborg et al 2013^26^. Briefly, nerves were fixed in 0. 2M phosphate buffer containing 2.5% PFA and 1.5% glutaraldehyde O/N at 4°C. Nerves were further incubated in 1% osmium tetroxide/1.5% potassium ferricyanide in water for 90 minutes at RT. After 3 x 5 minute washes in water, nerves were incubated in 1% tannic acid in 0.05M sodium cacodylate buffer for 2 hours at RT twice. Nerves were then washed in water again and osmicated in 1% osmium tetroxide for 30 minutes at RT. Nerves were then incubated in uranyl acetate, dehydrated and embedded in epoxy resin as described above. Following this, 1-2 mm thick cross sections were cut from the resin block, mounted with cyanoacrylate glue onto a specimen pin and 70 nm thick sections were examined by TEM to identify the regions of interest. Samples on pins were then carbon coated and mounted in the 3View microtome (Gatan). Once aligned, the sample and microtome were returned to the SEM chamber and put under vacuum. The regions of interest on the block face were re-located in the SEM using backscattered electron detection and the imaging and cutting parameters were optimised for each sample. Data sets were collected with section thickness between 120 and 150 microns in a Zeiss Sigma FEG-SEM coupled to Gatan 3View. Data was imported into Amira software (Thermo Fisher Scientific), where the cells of interest were manually segmented, reconstructed and rendered in 3D.

### Assessment of BNB function by Evans Blue

This experiment was performed as described in Napoli et al 2012^11^. Briefly, nerves were snap frozen in OCT upon dissection. Sections of 10μm were obtained using a cryostat, and mounted under a glass coverslip with fluoromount immediately. The nerves were immediately imaged with a Zeiss Axio imager microscope using the red channel.

### Assessment of the permeability of the perineurium

The permeability of the perineurium was assessed both *in situ* and *ex vivo*.

*Ex vivo*, Control nerves were harvested and immersed in a solution of 10% NaCl for 10 minutes. NaCl treated, Control or P0-RafTR samples were then immersed in a solution containing 40KDa dextran (10 mg/ml) for 15 min for permeability evaluation (being careful not to immerse the extremities).

*In vivo*, the animals were culled and their sciatic nerve exposed. As a positive control, a solution of 10% NaCl was applied to the sciatic nerve of Control animals for 10 minutes. A solution containing 40KDa dextran (10 mg/ml) was then applied on NaCl-treated, Control or P0-RafTR nerves for 15 minutes.

### Image analysis and Statistical analysis

2D confocal images were opened and processed using Fiji. Electron microscopy images were opened with Fiji and subjected to the A posteriori shading correction plugin developed by Maxime Pinchon, Laetitia Pasquet and Noël Bonnet and available online (automatic mode, 2x2). 3D Iba1 and CD31 immunostaining, as well as the BSA-FITC images were deconvoluted using the Huygens software and opened using Imaris.

The number of animals used in each experiment is stated throughout the paper. All the statistical analysis was performed using Prism (GraphPad). Before applying statistical test, all datasets were subjected to a Shapiro-Wilk normality test to determine the choice of the test. In cases when the normality of all groups was achieved, we performed a standard t-test to compare 2 conditions (Fig2, Fig 4, Ext.Fig.4, Ext.Fig.7, Ext. Fig. 8, Ext. Fig. 9d). When the normality was not achieved, we applied a Mann-Whitney test (Ext. Fig.9f) to compare the 2 conditions. For the assessment of the permeability of the perineurium, 3 groups were compared with a 1-way ANOVA, followed by a Bonferonni post-test (Ext. Fig.6). * is p-value <0.05, ** is p-value <0.01, *** is p-value < 0.001, **** is p-value <0.0001.

**Supplementary Information** is available in the online version of the paper

## Acknowledgements

This study was supported by programme grant from CRUK (C378), MRC project grant, MR/N009169/1, a Wellcome PhD studentship awarded to LM and MRC funding to the MRC LMCB University Unit at UCL, award code MC_U12266B. We would like to thank UCL Biological Services and Plexxikon Inc. for the provision of the PLX5622 formulated chow.

## Author contributions

L.M., I.N. and A.C.L. conceived and designed the project. L.M., I.N., I. J.W, S.S, A.B. performed experiments. L.M and A.C.L. wrote the manuscript. All authors reviewed the manuscript.

## Author information

The authors declare no competing financial interests. Correspondence and requests for materials should be addressed to A.C.L.

---

**Extended Data Figure 1.**
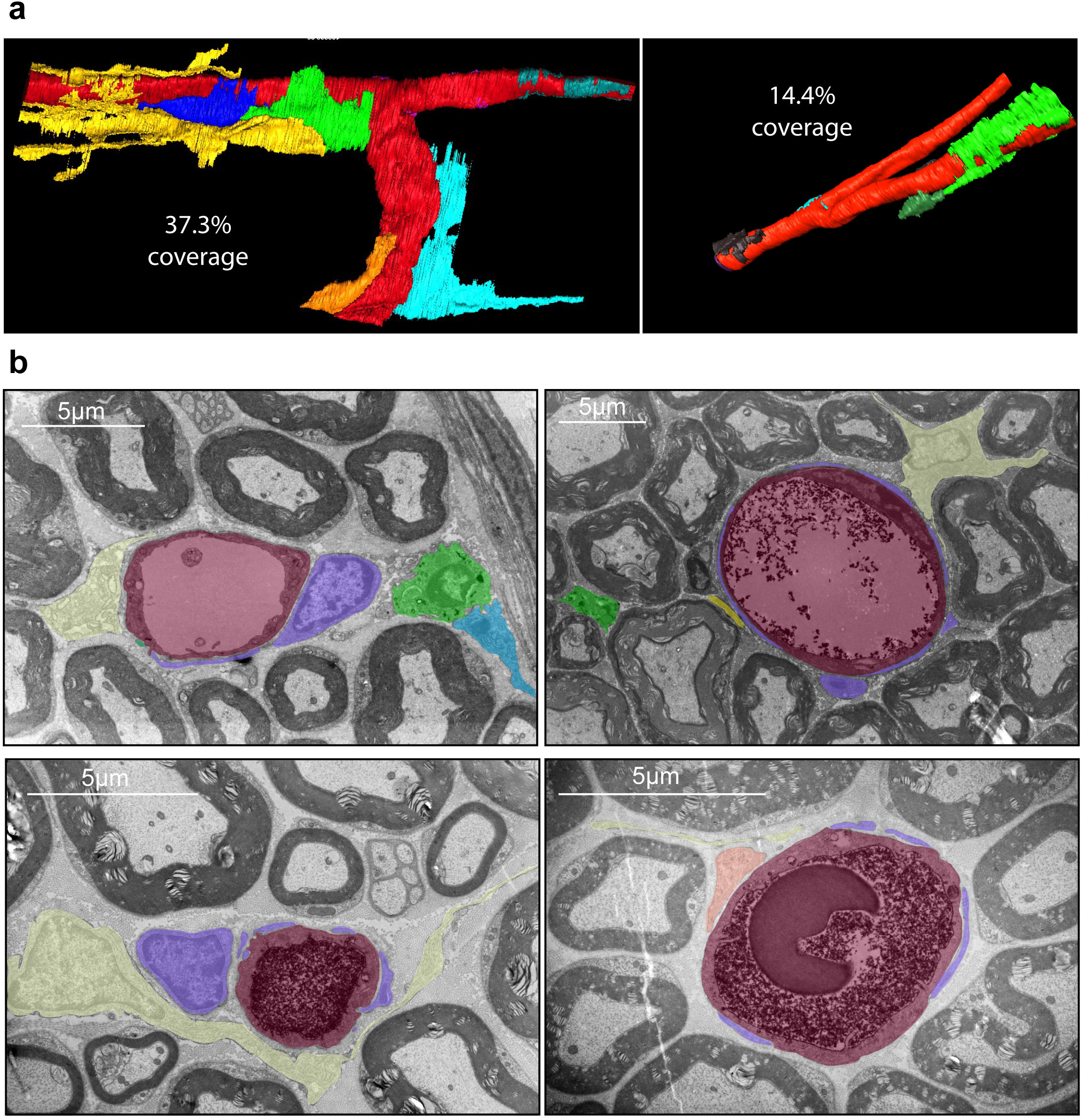
Incomplete coverage of endothelial cells in peripheral nerve. **a**, 3D EM reconstructions of blood vessels from the endoneurium of sciatic nerve. The ECs are coloured red with interacting cells individually coloured. The percentage coverage is indicated for each vessel. **b**, Representative 2D EM images of endoneurial blood vessels showing cellular interactions with the ECs. The endothelial cells and lumen are coloured magenta, with the pericytes within the basal lamina coloured purple. Other interacting cells are individually coloured. Note that while the pericytes are tightly associated with the blood vessels, other cell types are making looser contacts. Moreover the degree of coverage varies between the vessels.

**Extended Data Figure 2.**
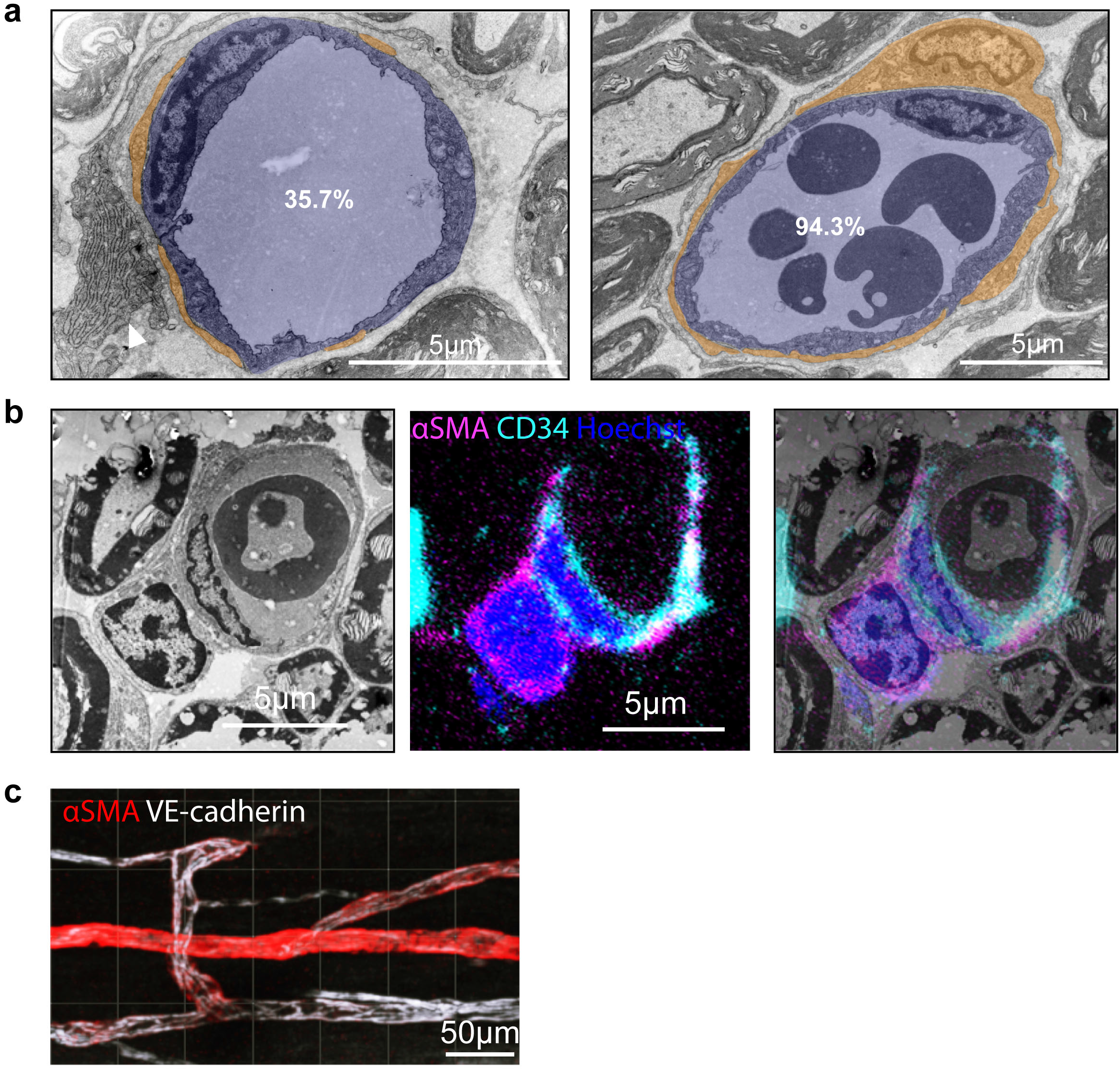
Pericyte coverage is highly variable between vessels. **a**, Representative EM images of endoneurial blood vessels showing the variability in pericyte coverage. The ECs and lumen are coloured blue whereas the pericytes are coloured brown and are identified by their localisation within the basal lamina. The white arrowhead indicates another cell type, a pericyte-like cell, which can be identified by the high levels of endoplasmic reticulum and extended thin protrusions that make multiple contacts with the basal lamina of the vessel. **b**, Pericytes in sciatic nerve are the only αSma+ cell in the endoneurium^13^, CLEM analysis confirms the localisation of αSma+ cells within the basal lamina and that they are closely associated with the ECs. **c**, 3D image of blood vessels within the endoneurium of an PACT cleared sciatic nerve showing variable coverage of ECs labelled for VE-cadherin (white) with αSma+ (red) pericytes.

**Extended Data Figure 3.**
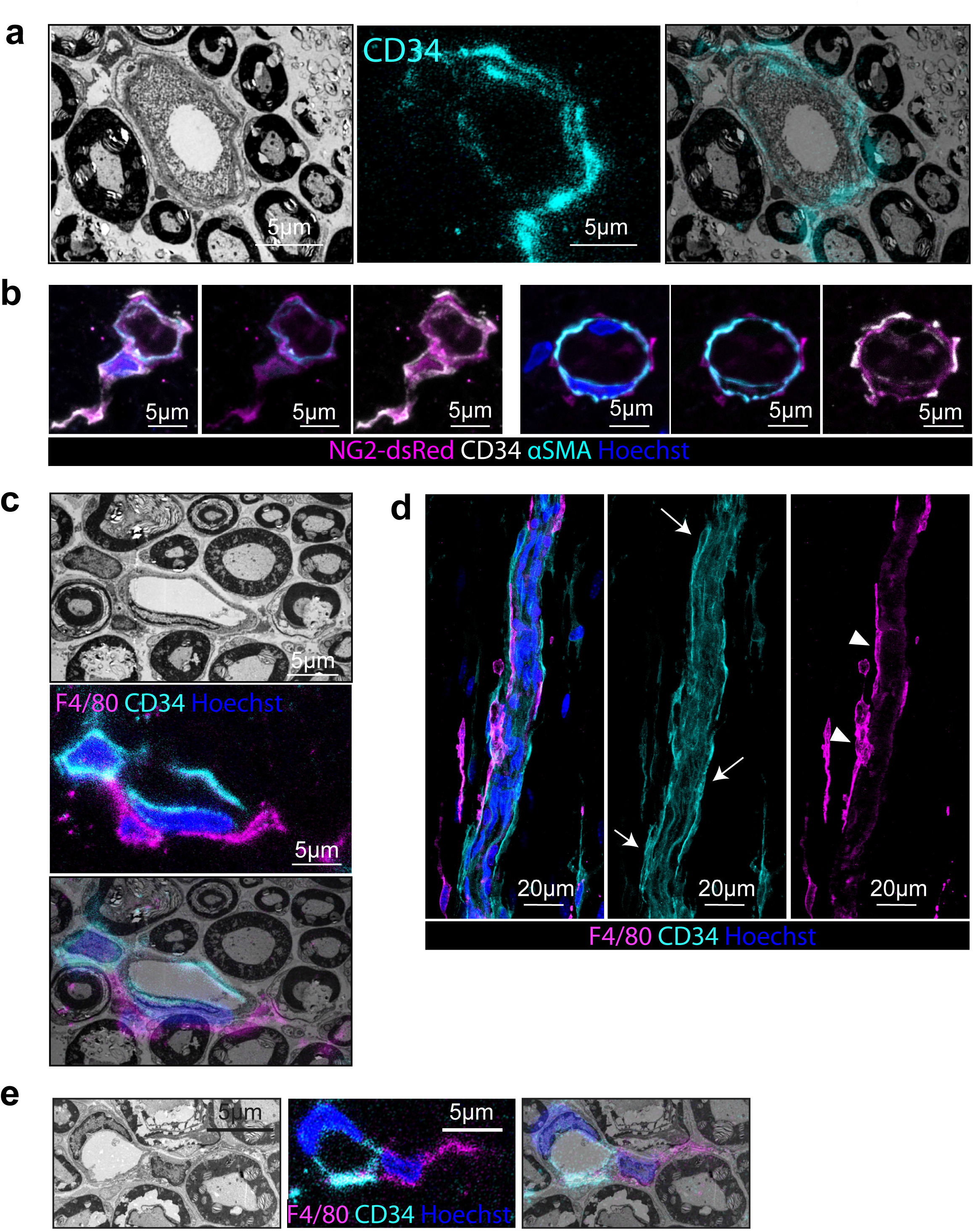
Macrophages and pericyte-like cells are closely associated with endoneurial blood vessels. **a**, Representative CLEM image shows that CD34+ cells are elongated cells, which interact with the basal lamina of blood vessels. **b**, We have previously identified a population of pericyte-like cells that are αSMA-/NG2+/PDGFRβ+/p75+ that make up 12.5% of cells within the endoneurium of which ~40% are loosely associated with blood vessels^13^. Here, representative confocal images show that these NG2+ (magenta), αSMA-(cyan) cells are also positive for CD34 (white). In contrast, the classical αSMA+ pericytes are negative for CD34. **c**, Representative CLEM image showing a macrophage (magenta) and a pericyte-like cell closely associated with a blood vessel. Arrows point to pericyte-like cells making close contacts along the vessel, arrowheads indicate perivascular macrophages. **d**, Confocal images of longitudinal sections of sciatic nerve immunostained to detect macrophages (F4/80, magenta), ECs (CD34, cyan) and pericyte like-cells (CD34, cyan), nuclei are labelled with Hoechst. **e**, Representative CLEM image showing a F4/80 positive macrophages (magenta) closely associated with endoneurial blood vessels (CD34, cyan). Prior work shows that resident F480/Iba1+ macrophages are 8% of cells within the endoneurium^13^.

**Extended Data Figure 4.**
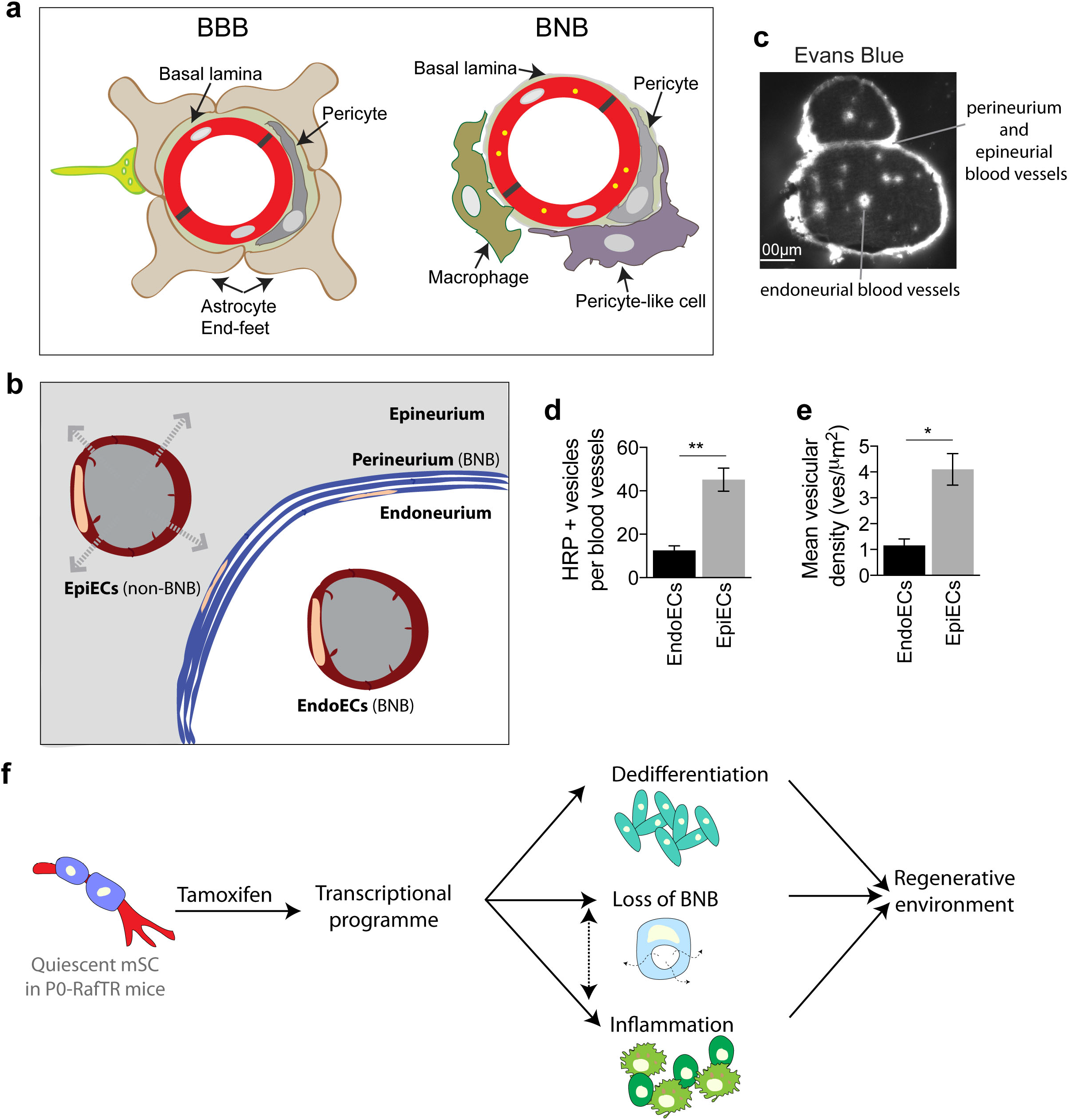
The BBB is distinct from the BNB. **a**, Cartoons representing the distinct cellular compositions that constitute the neurovascular unit of the BBB and the vascular unit of the BNB. Note that the ECs of the CNS are completely covered by pericytes and the end-feet of astrocytes. Astrocytes do not exist in the PNS and the vessels of the PNS are incompletely covered by closely associated pericytes and more loosely associated macrophages and pericyte-like cells. **b**, Cartoon illustrating the two components of the BNB; the perineurium that encloses the endoneurium of each nerve fascicle and the blood vessels within the endoneurium. In contrast, the blood vessels between the fascicles of the nerve are without barrier function. The grey shading shows the area that is penetrated by tracer following IV injection. **c**, Fluorescent image showing a transverse section of a sciatic nerve from an animal injected IV with the tracer Evans Blue. The dye (white) remains in the blood vessels of the endoneurium, whereas leakage is observed around the fascicles with entry into the endoneurium prevented by the perineurium. **d-e**, Quantification of transcytosis levels in endoneurial endothelial cells (EndoECs) and epineurial blood vessels (EpiECs) following the IV injection of HRP. Nerves were harvested 5 minutes after the injection, were processed for EM and the number of HRP+ vesicles were counted (n= 3 animals). Graphs show **d**, the number of HRP+ vesicles per blood vessel (mean±SEM). **e**, the mean vesicular density (number of HRP+ vesicles/ area of cytoplasm (mean±SEM). **f**, Cartoon representing the P0-RafTR transgenic mouse^11^ in which activation of Raf-kinase in mSCs by the injection of tamoxifen mimics an injury response and leads to the reversible opening of the BNB.

**Extended Data Figure 5.**
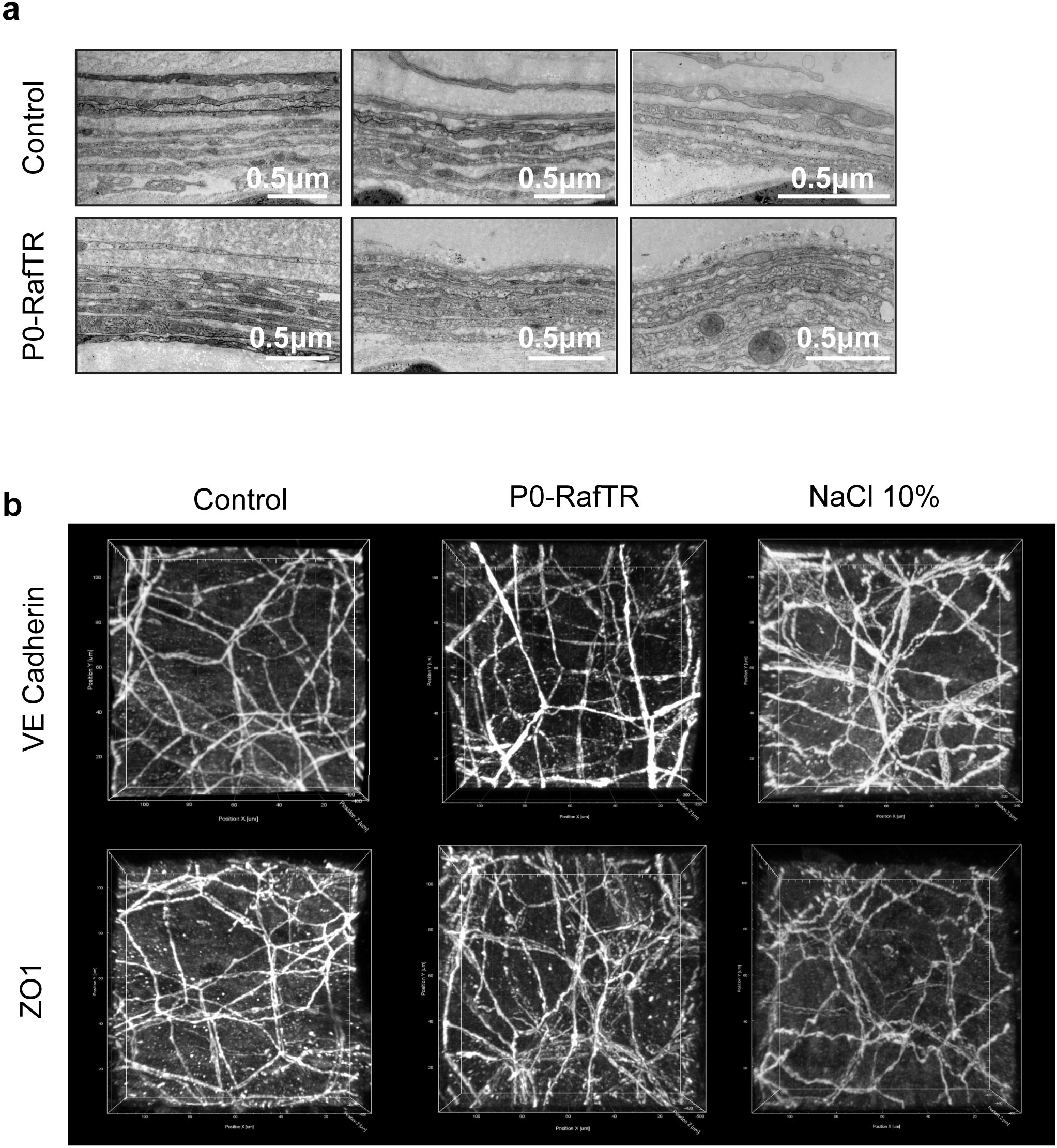
The structure of the perineurium remains similar when the BNB is open. **a**, Representative EM images of the perineurium in Control and P0-RafTR animals at Day 10 following the injection of tamoxifen. The structure of the perineurium appears similar in both groups of animals. **b**, Representative fluorescence images of junctional staining (VE cadherin/ZO1) of the perineurium in Control and P0-RafTR animals at Day 10 following the injection of tamoxifen and Control nerves treated with 10% NaCl for 15 mins to disrupt the barrier of the perineurium^27^. Note the junctional staining is similar between the Control and P0-RafTR animals whereas VE cadherin staining is more diffuse and the ZO1 levels appear lower in the NaCl treated nerves.

**Extended Data Figure 6.**
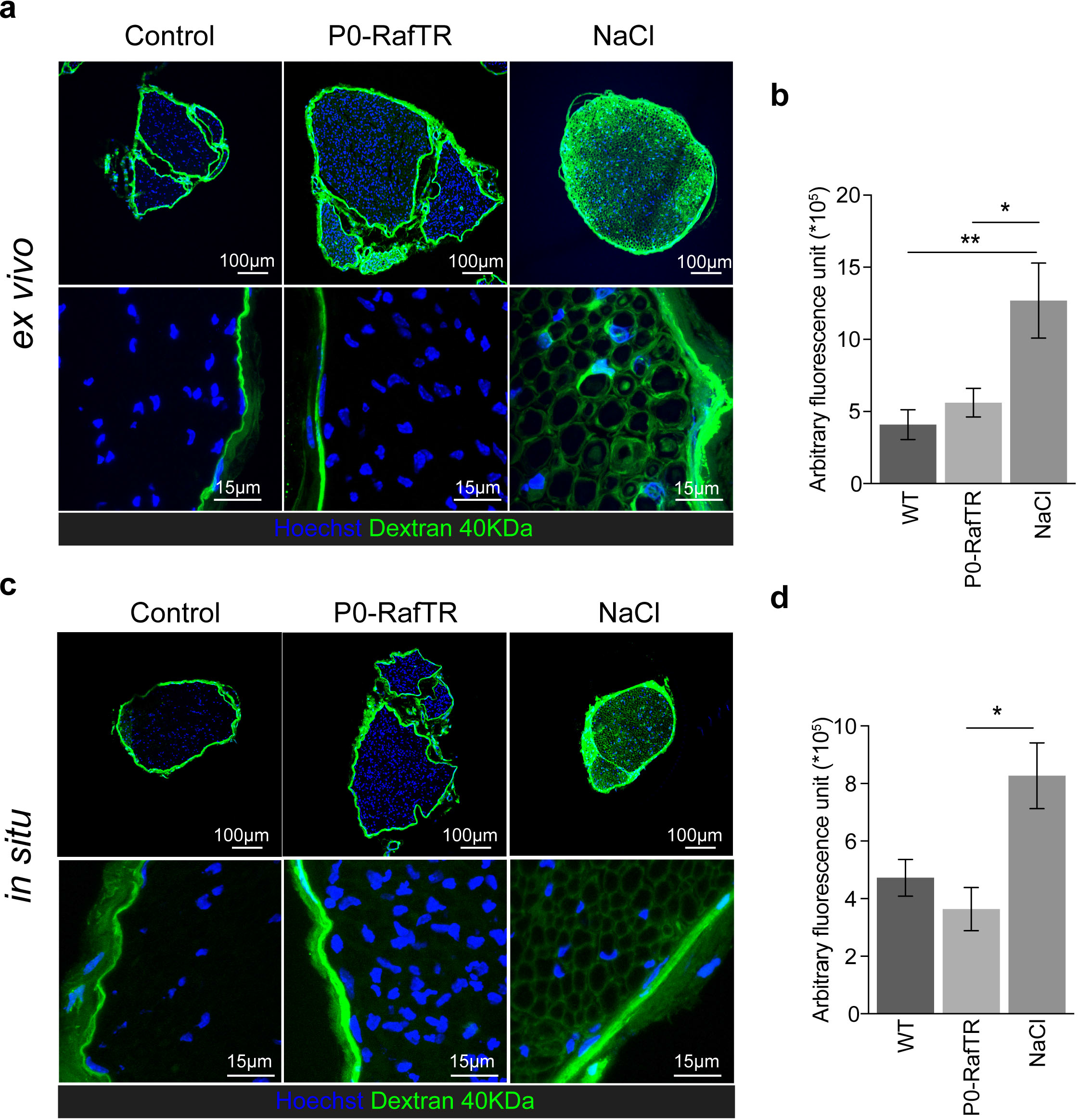
The barrier function of the perineurium is not compromised when the BNB is open. **a**, Representative images of transverse sections of sciatic nerves from Control and P0-RafTR nerves at Day 10 following tamoxifen treatment and Control nerves treated with 10% NaCl. The isolated nerves were incubated in a solution of 40KDa Dextran for 15 minutes after harvesting. Note, the tracer (green) only permeates into the endoneurium of the NaCl-treated nerves. Nuclei are labelled with Hoechst. **b**, Quantification of **(a)** showing the intensity of the green fluorescent signal in the endoneurium of Control, P0-RafTR and Control NaCl-treated nerves (n= 5, 6,and 4 animals respectively). **c**, As in **(a)** but the nerves were instead exposed to a solution of 40KDa Dextran in live anesthetised animals for 15 minutes prior to harvesting. Note that as in **(a)**, the dye only permeates into the endoneurium of the NaCl-treated nerves **d**, Quantification of **(c)** (n= 5, 4 and 2 animals respectively).

**Extended Data Figure 7.**
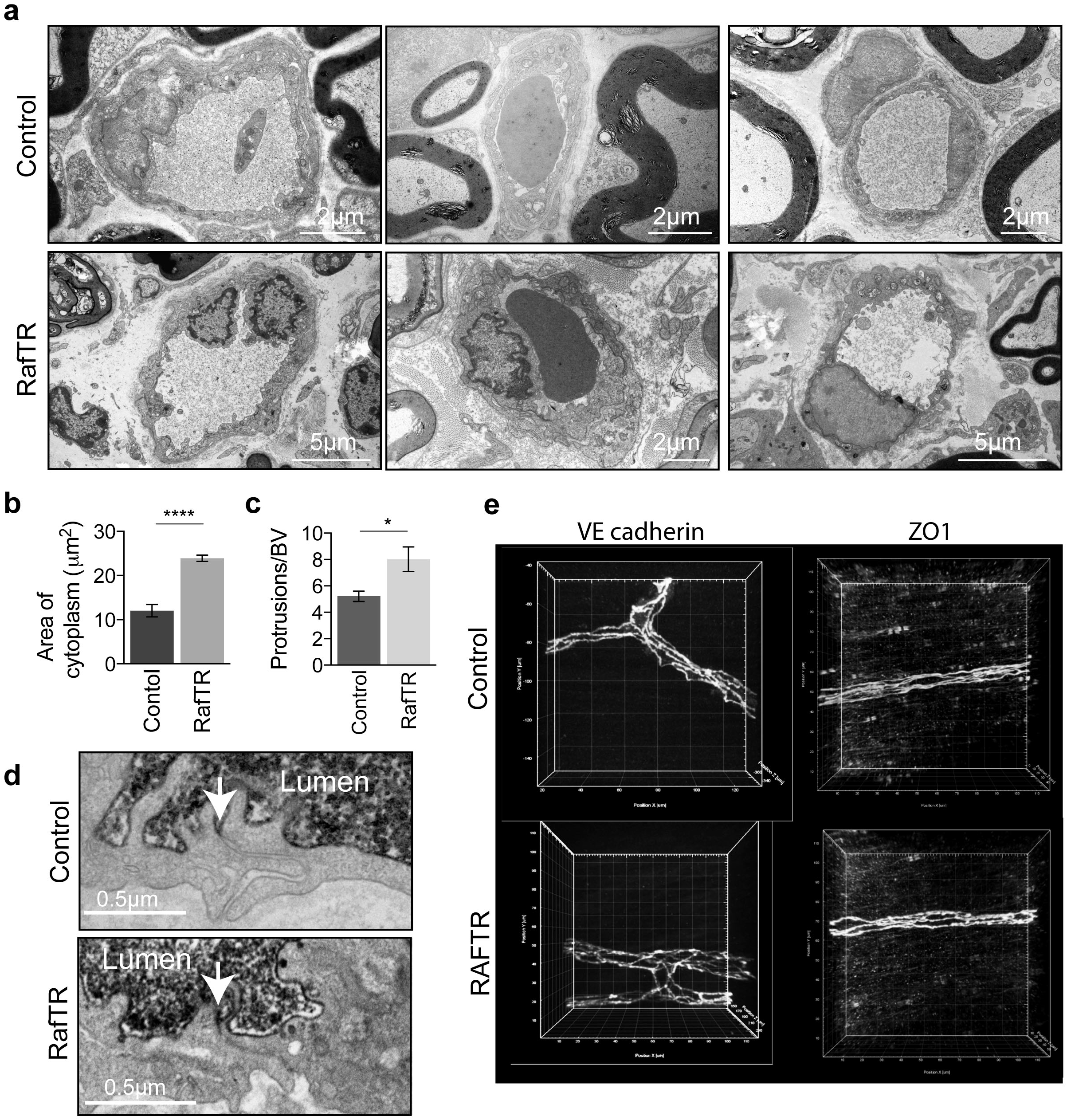
Endoneurial blood vessels show intact junctions when the BNB is open. **a**, Representative EM images of endoneurial blood vessels in Control and P0-RafTR animals at Day 10 following the injection of tamoxifen when the BNB is open in the P0-RafTR mice. Note the vessels have a more compact structure in the Control compared to the P0-RafTR animals. **b**, Quantification of the cytoplasmic area of endoneurial blood vessels in Control and P0-RAFTR animals (mean±SEM, n= 3 /4 animals). **c**, Quantification of the number of abluminal protrusions per endoneurial blood vessel in Controls and P0-RafTR animals (mean± SEM, n= 6 animals). **d**, Representative EM images of Control and P0-RafTR animals treated with tamoxifen for 10 days and injected with HRP. The penetration of HRP into the junctions is marked by arrows. Note the HRP does not penetrate into the junctions. **e**, Representative confocal images of cleared sciatic nerves immunostained for the junctional proteins VE-cadherin and ZO1, and in Control and P0-RafTR animals at Day 10 following tamoxifen injection. Note the junctional staining appears similar in the Control and P0-RafTR animals.

**Extended Data Figure 8.**
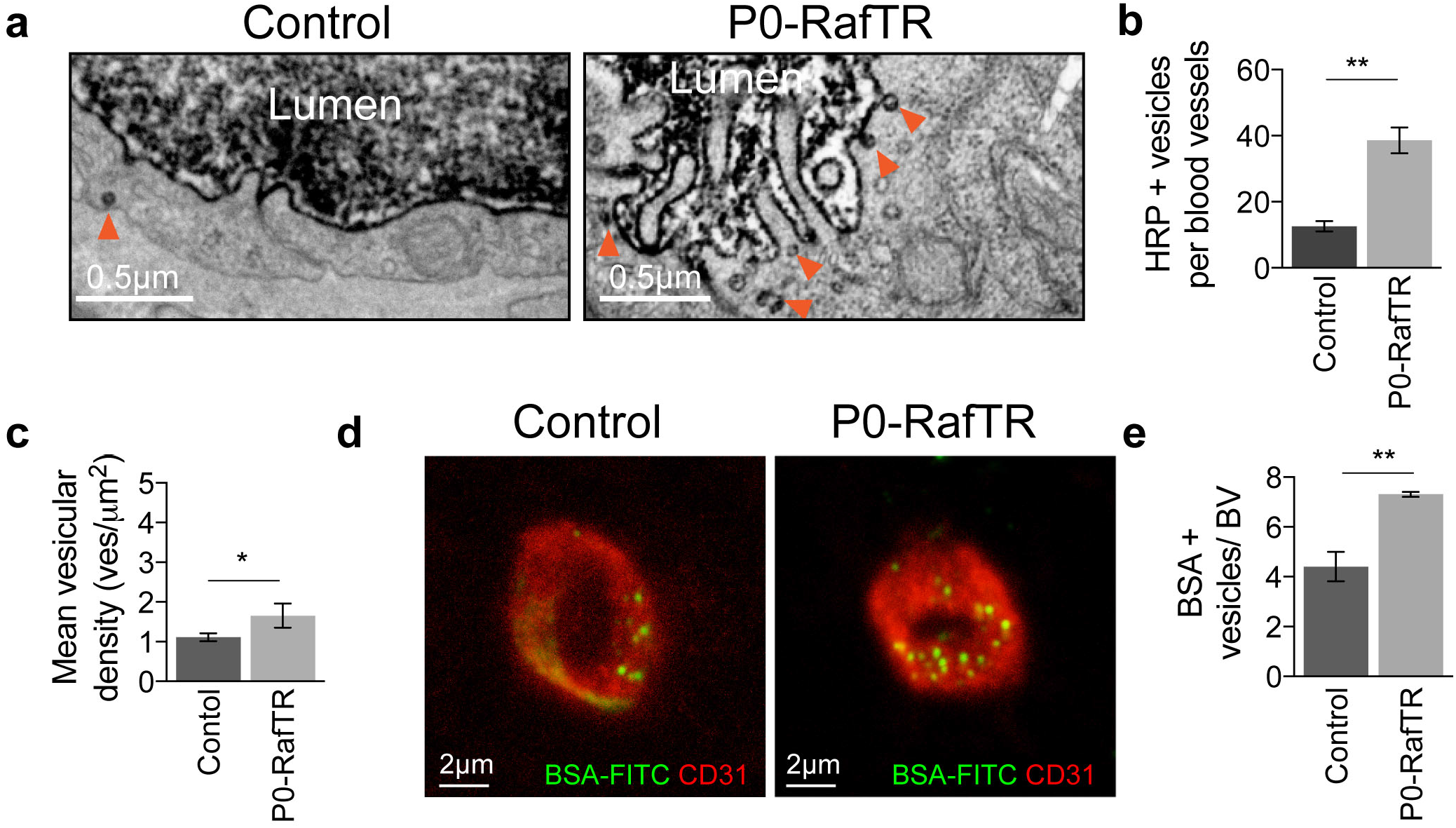
Endoneurial blood vessels have increased transcytosis rates when the BNB is open. **a**, Representative EM images of Control and P0-RafTR animals treated with tamoxifen for 10 days and injected with HRP. HRP+ vesicles are marked by arrowheads. Note the increased number of vesicles in the P0-RafTR animals. **b-c**, Quantification of transcytosis rates represented in **(a)**, graphs show b, the number of HRP+ per blood vessel (mean ± SEM (n= 3 -4 animals, t-test p-value= 0.0028), **c**, the mean vesicular density (mean± SEM (n= 3 -4 animals). T-test p-value= 0.0331) from Control and P0-RafTR animals at Day 10. **d**, Representative confocal images of endoneurial blood vessels from Control and P0-RafTR (Day 10) animals injected with BSA-FITC and counterstained with CD31. BSA-FITC+ vesicles are visible in the endothelial cells. **e**, Quantification of (d) (mean±SEM, n=4 animals).

**Extended Data Figure 9.**
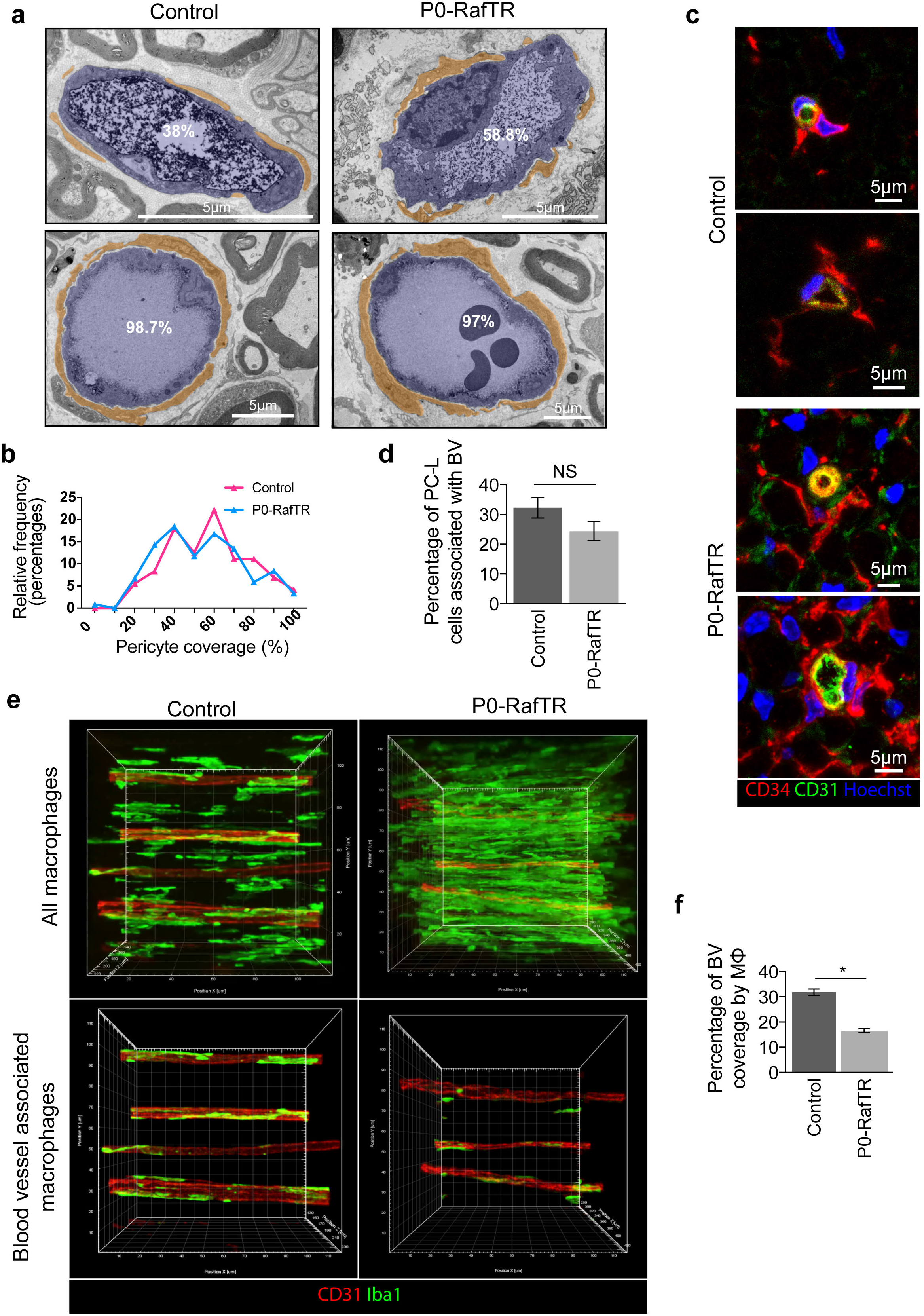
Macrophages move away from blood vessels when the BNB is open. **a**, Representative EM images showing examples of endoneurial blood vessels in Control and P0-RafTR animals at Day 10 following tamoxifen injection that illustrate the variability in pericyte coverage in both genotypes. **b**, Quantification of **(a)** showing the relative frequencies of the percentage ECs coverage by pericytes (n=3/4 animals). **c**, Representative confocal images of transverse sections of sciatic nerve showing examples of pericyte-like cells (CD34+, red, CD31-, green) associated with endoneurial blood vessels identified by the labelling of ECs (CD34+, red, CD31+, green) in Control and P0-RafTR mice at Day 10 following treatment with tamoxifen. **d**, Quantification of **(c)** showing the percentage of pericyte-like cells (PC-L) that are associated with blood vessels. Note that pericyte-like cells remain associated with blood vessels when the barrier is open (mean±SEM, n= 4 animals per condition). **e**, Representative confocal images showing the localisation of macrophages in the endoneurium of sciatic nerves. Nerves from Control and P0-RafTR animals were harvested at day 10 following treatment with tamoxifen. Nerves were cleared using iDISCO techniques and were immunostained to detect macrophages (Iba1,green) and blood vessels (CD31, red). The upper images show all macrophages whereas the lower images have had the macrophages that are not associated with the blood vessels removed. The images show the close association of a population of macrophages with the blood vessels and that despite a large increase in macrophages in the P0-RafTR animals following tamoxifen treatment, there are fewer in the perivascular region.**f**, Quantification of **(e)** showing the percentage of blood vessels coverage by macrophages (mean±SEM, n= 4 animals).

**Extended Data Figure 10.**
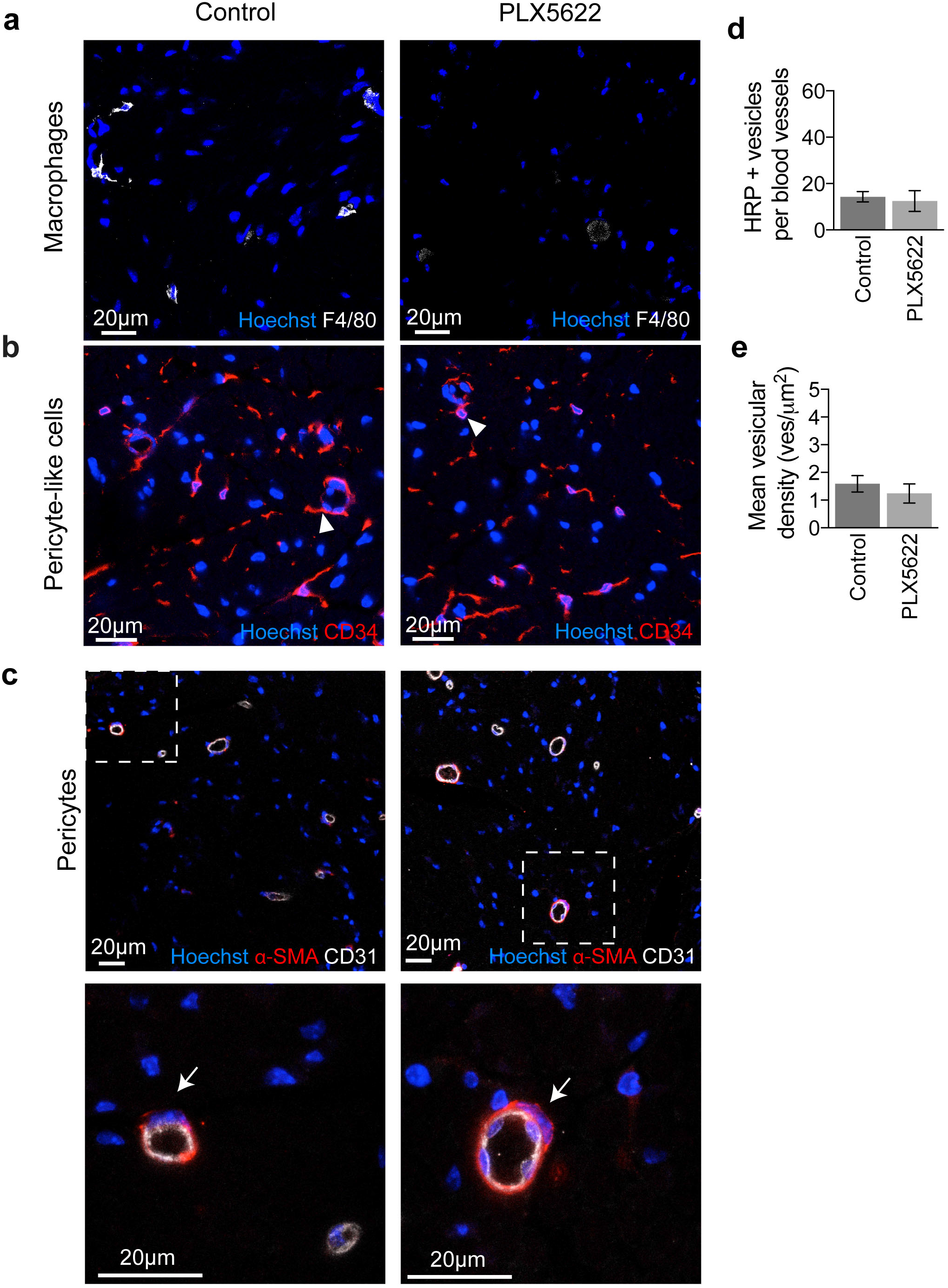
PLX5622 efficiently and specifically depletes macrophages from the nerve endoneurium. Representative confocal images of the endoneurium of Control and PLX5622 treated animals immunostained to detect **a**, macrophages (F4/80, white). **b**, pericyte-like cells (CD34, red). Arrowheads indicate pericyte-like cells in close association with blood vessels. **c**, pericytes (αSMA, red) and ECs (CD31, white). The lower panel depicts a higher magnification of the region shown in the upper panel. Arrows point to the nuclei of pericytes. Nuclei are counterstained with Hoechst (blue). **d-e**, Quantification of transcytosis rates in Control and PLX5622 treated animals (n= 5/6) from EM images as shown in Figure 4b. Graphs show **d**, Number of HRP+ vesicles per endoneurial blood vessel (mean±SEM), e, Vesicular density of endoneurial blood vessels (mean±SEM).

**Movie 1 Incomplete coverage of endothelial cells in peripheral nerve**.

3D EM reconstructions of a blood vessel from the endoneurium of sciatic nerve. The ECs are coloured red with interacting cells individually coloured.

**Movie 2 Macrophage interaction with endoneurial blood vessels**.

Representative confocal images showing the localisation of macrophages in the endoneurium of sciatic nerves. Nerves from Control and P0-RafTR animals were harvested at day 10 following treatment with tamoxifen. Nerves were cleared using iDISCO techniques and were immunostained to detect macrophages (Iba1,green) and blood vessels (CD31, red). The first sequence shows a representative nerve from a Control animal. The second sequence shows a representative nerve from a P0-RafTR animal.

